# Elasticity of the HIV-1 Core Facilitates Nuclear Entry and Infection

**DOI:** 10.1101/2023.09.29.560083

**Authors:** Akshay Deshpande, Alexander J. Bryer, Jonathan R. Andino-Moncada, Jiong Shi, Jun Hong, Cameron Torres, Shimon Harel, Ashwanth C. Francis, Juan R. Perilla, Christopher Aiken, Itay Rousso

## Abstract

HIV-1 infection requires passage of the viral core through the nuclear pore of the cell, a process that depends on functions of the viral capsid ^1,2^. Recent studies have shown that HIV-1 cores enter the nucleus prior to capsid disassembly ^3–5^. Interactions with the nuclear pore complex are necessary but not sufficient for nuclear entry, and the mechanism by which the viral core traverses the comparably sized nuclear pore is unknown. Here we show that the HIV-1 core is highly elastic and that this property is linked to nuclear entry and infectivity. Using atomic force microscopy-based approaches, we found that purified wild type cores rapidly returned to their normal conical morphology following a severe compression. Results from independently performed molecular dynamic simulations of the mature HIV-1 capsid also revealed its elastic property. Analysis of four HIV-1 capsid mutants that exhibit impaired nuclear entry revealed that the mutant viral cores are brittle. Suppressors of the mutants restored elasticity and rescued infectivity and nuclear entry. Elasticity was also reduced by treatment of cores with the capsid-targeting compound PF74 and the antiviral drug lenacapavir. Our results indicate that capsid elasticity is a fundamental property of the HIV-1 core that enables its passage through the nuclear pore complex, thereby facilitating infection. These results provide new insights into the mechanisms of HIV-1 nuclear entry and the antiviral mechanisms of HIV-1 capsid inhibitors.

---

The HIV-1 capsid, which forms the shell of the core and acts as a protective barrier, encapsidates the enzymes reverse transcriptase and integrase and the viral genomic RNA. During infection, the HIV-1 genome is transported into the nucleus and integrates into host cell chromosomal DNA. Recent studies suggest that the core remains fully or mostly intact during nuclear entry, thereby ensuring completion of reverse transcription within the nucleus ^3,4^. It is presently unclear how the core passes through the nuclear pore ^6–8^ while remaining intact. Docking of the HIV-1 core at the NPC and subsequent nuclear entry are effected by interactions between the capsid and cellular factors, including cyclophilin A, CPSF6, and nucleoporins (e.g. Nup358 and Nup153) ^6,9–16^. Treatment with low concentrations of the capsid-binding compounds PF-3450074 (PF74) ^17^ and Lenacapavir (LCV) ^18^ reduces HIV-1 infectivity by inhibiting nuclear entry ^19–27^. Nuclear entry can also be reduced by mutations in the capsid protein (CA). For example, the capsid-stabilizing CA mutation E45A markedly reduces nuclear entry and consequently infectivity ^28,29^ (Fig. 1 A-D and Fig. 2). However, position 45 in CA resides at an inter-subunit interface in the capsid that is not a binding site for known host proteins that promote nuclear entry ^2^. In addition, E45 lies outside the binding site for PF74 and LCV, yet the mutant virus is resistant to these compounds ^20,30^. Remarkably, the suppressor mutation R132T rescues nuclear entry and infectivity of the E45A mutant and restores its sensitivity to PF74 without reversing the hyperstability of the capsid ^29^.

**Fig. 1:**
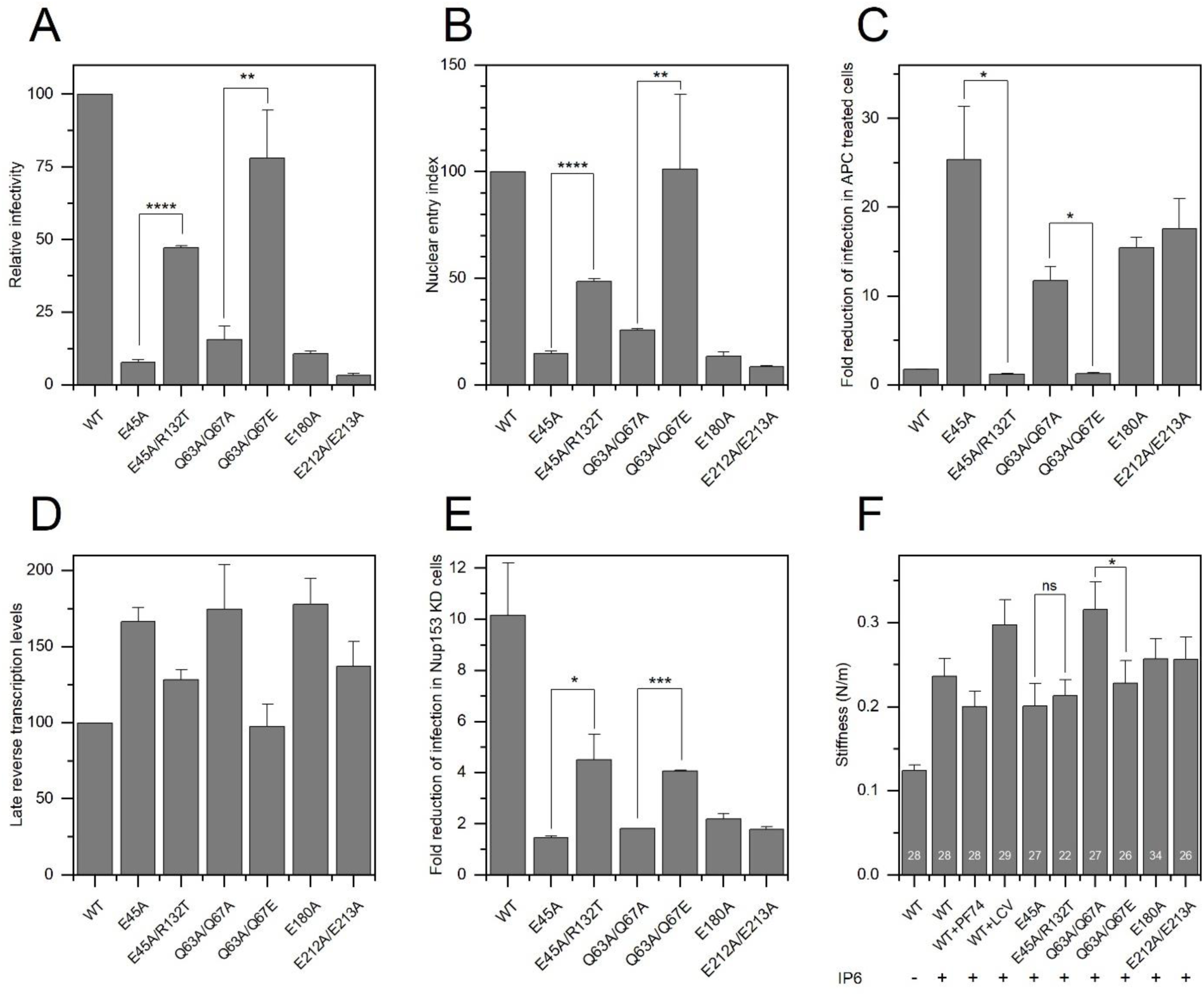
DNA synthesis, nuclear entry, infectivity and averaged measured point stiffness of isolated HIV-1 cores. (A) Relative single-cycle infectivity of wild type and CA mutant viruses. (B) Nuclear entry efficiency of CA mutants relative to WT HIV-1. Value represent the ratio of 2-LTR circle levels to late reverse transcripts. (C) Ratio of infection of control Hela cells vs. aphidicolin-treated cells by WT and mutant viruses. (D) Relative levels of late reverse transcripts in target cells. (E) Fold reduction of infection in Nup153-depleted cells vs. control cells. For A-E, values shown are the mean of three independent experiments, with error bars representing the standard error of the mean. (F) Stiffness measurements of purified HIV-1 cores. Each stiffness value was calculated as the average of ∼480 force–distance curves obtained from individual cores. Measurements were conducted in the presence of inositol hexakisphosphate (IP6; 100 μM) except for the first column. The t-test analysis revealed that differences between the stiffness values of the various samples and WT are statistically significant (p values: <0.001 and <0.05 for WT+IP6+PF74 and E45A+IP6, respectively, <0.0001 for the remaining samples). Error bars in all panels represent the standard error of the mean.

**Fig. 2:**
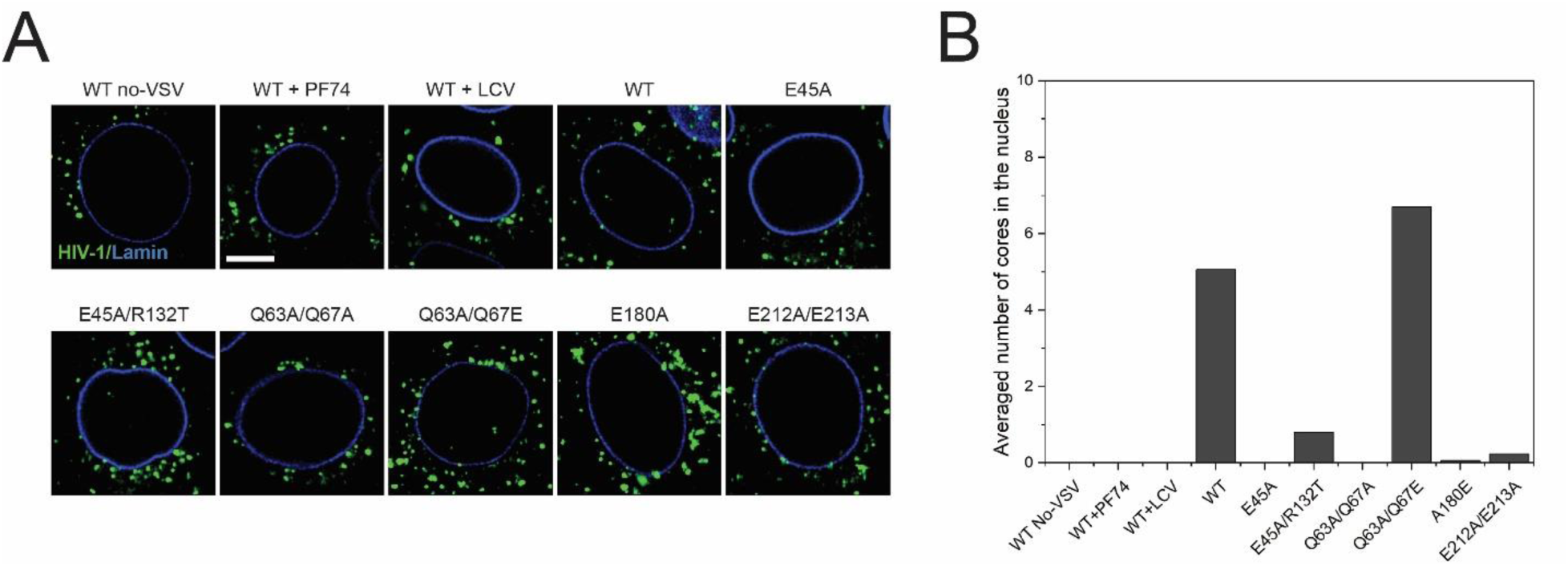
Fluorescence microscopy imaging of HIV-1 nuclear entry. (A) Representative single z-stack images and (B) quantification of HIV-1 INmNG puncta nuclear entry in TZM-bl cells acquired at 8 hours post-infection. The number of fluorescent INmNG puncta (fHIV-1, green) inside 0.5 um of the nuclear membrane (Lamin, blue) was scored from 30 nuclei in individual experiment from each of 3 independent experiments. Scale in (A) is 5 um.

In earlier work, we employed atomic force microscopy (AFM) to show that CA mutations can affect the physical property of the capsid known as stiffness, defined as the resistance to an applied force, To determine whether capsid stiffness plays a role in HIV-1 nuclear entry and sensitivity to capsid inhibitors, we quantified the stiffness of purified HIV-1 cores using AFM operated in the nano-indentation mode (Fig. 1F). Untreated WT cores exhibited a mean stiffness value of 0.12 ± 0.01 N/m, and addition of the capsid-stabilizing host cell metabolite inositol hexakisphosphate (IP6) ^31^ (100 μM) increased the stiffness to a mean value of 0.24 ± 0.02 N/m. Cores treated with both IP6 and the antiviral compounds PF74 (1.25 μM) or IP6 and LCV (500 pM) exhibited a comparable increase in stiffness (value of 0.20 ± 0.02 N/m or 0.30 ± 0.03 N/m, respectively). Thus, addition of the capsid-targeting antiviral compounds at concentrations that inhibit HIV-1 nuclear entry did not further increase the stiffness of HIV-1 cores that were treated with IP6. We analyzed three additional CA mutants Q63A/Q67A, E212A/E213A and E180A. Like E45A, these mutants exhibited low infectivity, impaired nuclear entry, and cell-cycle-dependent infection yet underwent efficient reverse transcription in target cells (Fig. 1A-D and Fig. 2). Moreover, infection by these mutants was minimally affected by depletion of Nup153 (Fig. 1E) ^32,33^. Purified cores from these mutants also exhibited increased stiffness. Q63A/Q67A cores showed the highest mean stiffness value of 0.31 ± 0.03 N/m, whereas the value for E180A and E212A/E213A cores was 0.26 ± 0.03 N/m. The pseudorevertant Q63A/Q67E emerged during prolonged culture of the Q63A/Q67A mutant in T cells and exhibited nuclear entry and infectivity similar to the wild type (Fig. 1 and 2). The mean stiffness value of this mutant was reduced relative to its parent mutant Q63A/Q67A (0.23 ± 0.03 vs. 0.31 ± 0.03, respectively; P= 0.0455). As previously reported, the suppressor mutation R132T partially rescued E45A infectivity and nuclear entry (Fig. 1A, B and Fig. 2), though it does not restore normal capsid stability ^29^. These variants exhibited similar mean stiffness values (0.20 ± 0.03 N/m for E45A and 0.21 ± 0.02 N/m for E45A/R132T; p > 0.05). Collectively, these observations indicate that rescue of E45A and Q63A/Q67A nuclear entry by R132T and Q67E, respectively, does not involve restoration of normal core stiffness, suggesting that another property of the core promotes nuclear entry and infectivity.

Studies of the HIV-1 capsid have demonstrated that the inter-subunit interfaces are structurally variable, indicating that the capsid is flexible ^34,35^. We hypothesized that capsid elasticity is required for HIV-1 nuclear entry. To quantify elasticity, we exploited the ability of the AFM probe to induce structural deformation in the viral core and analyzed the consequences on the mechanical behavior and structural integrity of the capsid. For this purpose, the AFM was operated in the nano-indentation mode but with high maximal loading forces (≥ 5 nN, vs. the 1–1.5 nN range used in stiffness measurements) to explore the mechanical tolerance limits of the cores and quantify their critical loading force: the force at which there is an abrupt drop in the force-distance curve, representing the failure and collapse of the structure ^36^. We observed no evidence for structural failure in the force–distance curves, even at a maximal loading force of 20 nN (Fig. 3A). Morphological analysis by scanning AFM further demonstrated that all cores (n=72) showed no evidence for structural damage following perturbation (Extended Data Fig. 1). For one core in the 10 nN applied force group, we observed a drop in the force–distance curve (Fig. 3A inset). Morphological analysis of this core structure showed that it remained intact and non-deformed. Therefore, this core likely underwent reversible buckling rather than breakage. The results show that HIV-1 cores can withstand marked deformation without undergoing irreversible structural damage, indicating that the HIV-1 capsid is highly elastic. This finding is in sharp contrast to other viruses, such as bacteriophage HK97, hepatitis B virus, and herpes simplex virus, which have brittle capsids that undergo structural failure at forces as low as 1-6 nN ^37^.

**Fig. 3:**
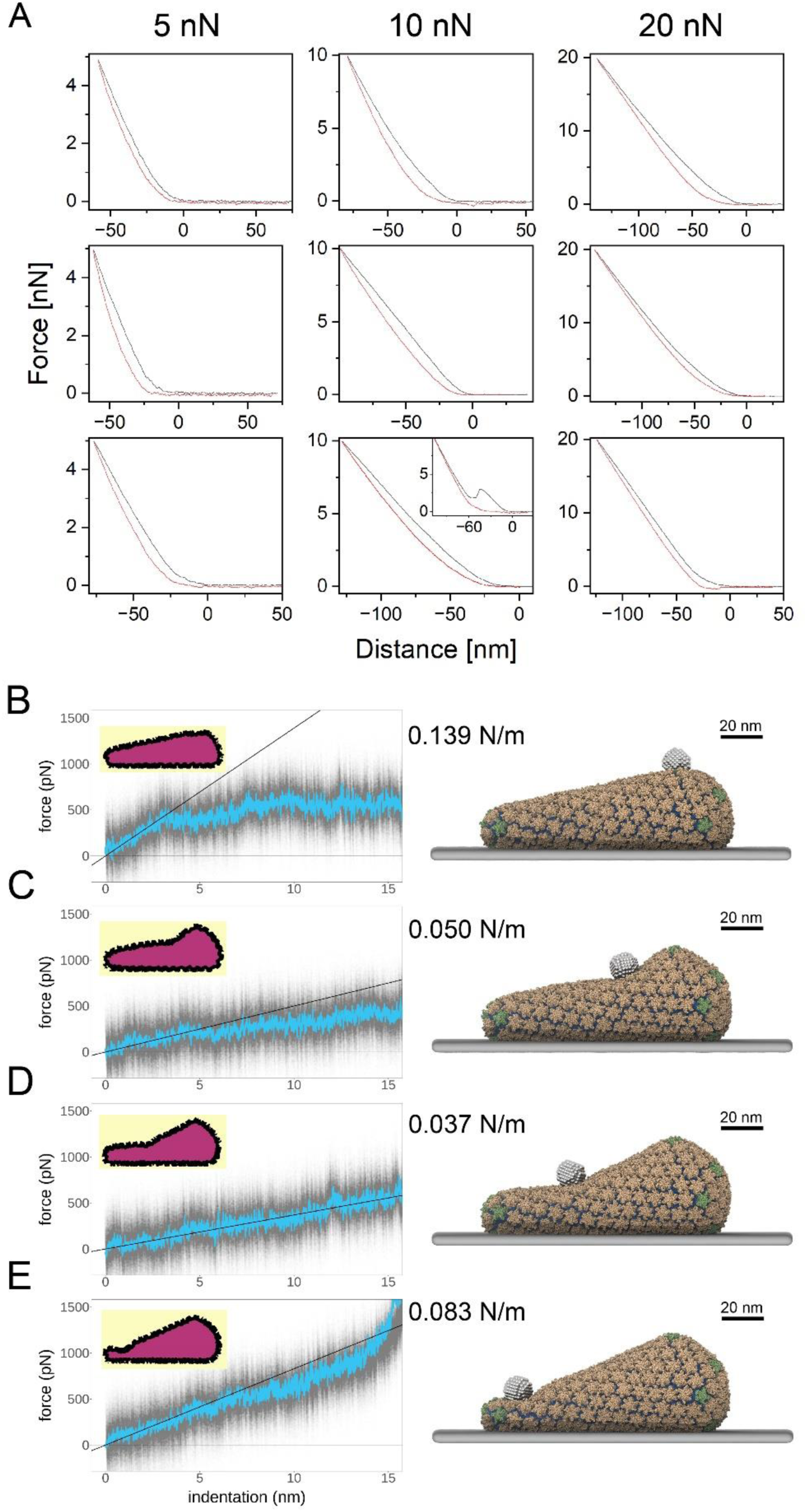
Experimental and simulated AFM nanoindentation of wild type IP6-treated cores. (A) Representative force–distance (FD) curves where force loading (black) and unloading (red) curves are shown. The maximal loading force was set to 5 nN (*n*=23), 10 nN (*n*=24), and 20 nN (*n*=25). Despite the application of high maximal loading forces, none of the curves exhibited any indication of structural failure in the core. In one core (of the 72 examined), we observed an abrupt drop in the loading curve (inset). (B-E) Simulated FD curves for nanoindentation of coarse-grained HIV-1 capsids, testing the effect of probe location, ultimately capsid curvature, on measured material properties. The plots show raw data in gray, and a windowed-average in blue. The inset volume cross-sections show the deformed capsid morphologies and provide a sense of global deformation. The solid black line shows a linear fit of the FD curve for the first 3 nm of indentation, which was used to derive the stiffness values given to the right of each plot. Additionally, snapshots from these nanoindentation simulations are provided on the right, with all components visualized: spherical AFM probes, capsids, and baseplates. (B) Probe located at the large end of the conical capsid. Note the global deformation, narrowing, of the cone in the inset graphic. (C) and (D) show probe locations in the relatively flat, mid-regions of the capsid. (E) Shows nanoindentation of the narrow, most highly curved region of the capsid.

To complement the AFM measurements, we performed coarse grained *in silico* computational nanoindentation simulations at four distinct regions along the wild-type capsid surface to model the resistance of the capsid to deformation. After placing four spherical AFM probes equidistant along the capsid’s surface (Fig. 3 B-E), we modeled a constant pushing velocity to the tip (3.125 nm/μs). While the simulated tip velocity is higher than what is achievable in physical experiments, simulations can provide a high temporal and spatial resolution view of how the capsid deforms and responds to mechanical stress. Indentation of the broader region of the capsid initially showed the highest stiffness among the four probed regions, with the capsid appearing to buckle after 3 nm of indentation demonstrating a change in the elasticity of the capsid’s mechanical properties at indentation depths beyond 3 nm (Fig. 3B). Indentations in the relatively flat, middle regions of the capsid revealed the lowest stiffness (Fig. 3C, D) and the most linearity in FD-curves. The narrow end of the capsid showed high stiffness through the indentation profile, until contact with the baseplate is made (Fig. 3E). Our simulations support our experimental AFM data and show that the wild-type capsid is an elastic container, able to withstand deformations without failure of the intermolecular interfaces comprising the CA lattice. The different stiffness values observed at different positions further suggests that the HIV-1 capsid has a material heterogeneity that is determined by its variable curvature.

Simulated nanoindentation of E45A and E45A/R132T capsids showed that the mutations produced a significant increase in stiffness (Extended Data Fig. 2) and, additionally, affect capsid deformability (Extended Data Figs. 2-5), with E45A capsids in particular rupturing at relatively smaller indentation distances compared to wild type and E45A/R132T while absorbing more power through deformation. Importantly, when we considered simulated probe locations, and thus regions of the capsid, independently, we found that wild type, E45A and E45A/R132T capsids have similar stiffness in the low curvature mid-regions of the capsid (Extended Data Fig. 6). At the highly curved broad end, the wild type was slightly stiffer than either mutant, further demonstrating that the capsid’s material response is predicated by its shape and location of indentation. The simulations show that the wild-type capsid is a robust and elastic container, able to sustain extreme deformations and recover (Extended Data Fig. 7) prior to failure of the intermolecular interfaces comprising the CA lattice. The simulations further suggest that the capsid has a material heterogeneity that is dictated by its curvature.

To physically assess the elasticity of HIV-1 cores, we analyzed the shape of the viral core soon after release of strong applied force. In this approach, a core was first imaged by scanning at a low loading force of 300 pN (Figure 3A). We then selected a region of interest (Extended Data Fig. 8) located in the main body of the core and rescanned it. This region was then scanned at a high loading force (5 nN) that caused the core to compress dramatically at this region. The force was immediately reduced to 300 pN and the compressed region rapidly imaged repeatedly over time (Fig. 4B). The volume under the scanned region was then calculated as a function of time. Fig. 4C shows two representative trajectories in which the volume was compressed to 50% and 30% of the initial volume and recovered to nearly 100% and 80%, respectively. Mean values of the compressed volume (pressed: 39 ± 3%), the immediate recovery volume (Immediate: 92 ± 1%), and the volume 5–7 minutes after compressing the core (later: 92 ± 1%) were determined from analysis of 36 individual cores (Fig. 4D). Wild type HIV-1 cores displayed nearly complete recovery after being strongly compressed. The measured recovery volume was highly consistent individual core volume trajectories. Consequently, the averaged immediate and late volumes were identical, indicating that the core recovered its initial shape in a time shorter than the temporal resolution of the assay (10–15 seconds) (Fig. 4C and D). These results further demonstrate that HIV-1 cores are highly elastic and rapidly recover nearly their full initial volume after being strongly compressed. For the majority of cores, we observed no structural damage in the compressed region of the CA-lattice. In 14% of cores analyzed, (5 of 36), structural damage was apparent near the end of the core following compression and recovery (Fig. 5A). We previously reported that capsid disassembly begins at the end of isolated cores during *in vitro* reverse transcription. This highly curved and pentamer-rich section has been proposed to be the least stable region in the HIV-1 capsid ^38^. Our data are consistent with this model, in which the weakest region in the structure collapses as the core is compressed.

**Fig. 4:**
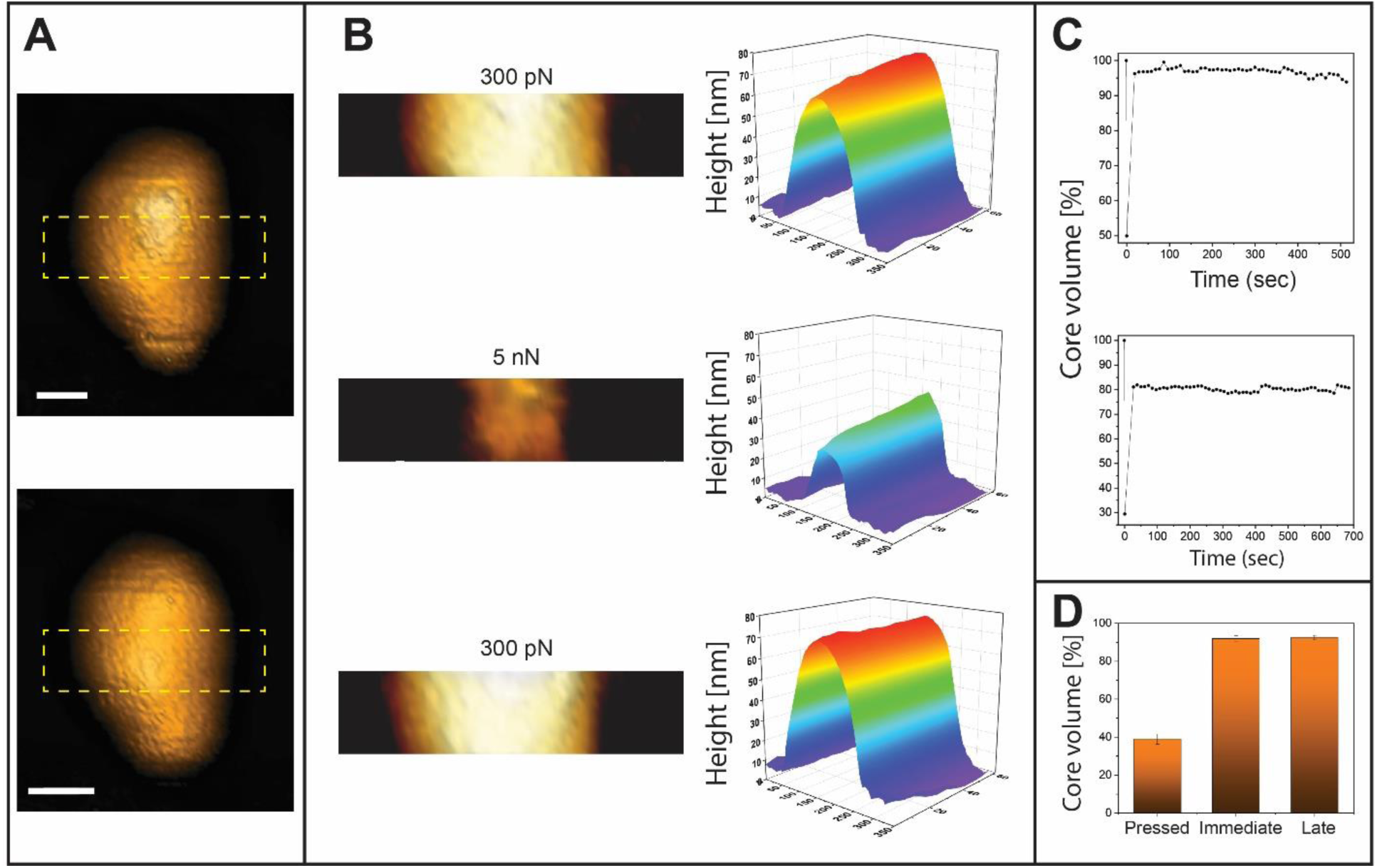
Volume recovery of the HIV core following compression. (A) Topographic AFM images of an isolated IP6-treated WT core before (upper image) and after (lower image) extreme compression. Both images were acquired using the QI mode at a maximal loading force of 300 pN. Subsequent measurements focused on a selected central region, as marked by a dashed yellow rectangle. Scale bars are 50 nm. (B) Representative topographic AFM images (left panels) and the corresponding Matlab plots (right panels) for the selected central region of an isolated IP6-treated WT core before (top), during (middle), and immediately after (bottom) compression by a 5 nN loading force. The same region was repeatedly scanned and imaged over time in QI mode at maximal loading force of 300 pN. (C) Two representative volume recovery trajectories (of the 36 trajectories obtained). The volume beneath the selected region was calculated using MatLab and plotted as a function of time. The initial volume was set as 100%, and all other measurements were normalized accordingly. The upper curve shows nearly full volume recovery, whereas the lower curve shows 80% recovery, which was the minimum volume recovery percentage we detected. (D) The average volume recovery of the 36 analyzed cores. The average compressed volume was 39% (pressed). The average volume upon reducing the force back to 300 pN was 92% (immediate). Several minutes (5–7 min) later, the recovery volume remained unchanged at 92% (late). T-test analysis revealed that the difference between the compressed and immediate or late recovered volume is significant (p value <0.0001). Error bars represent the standard error of the mean.

**Fig. 5:**
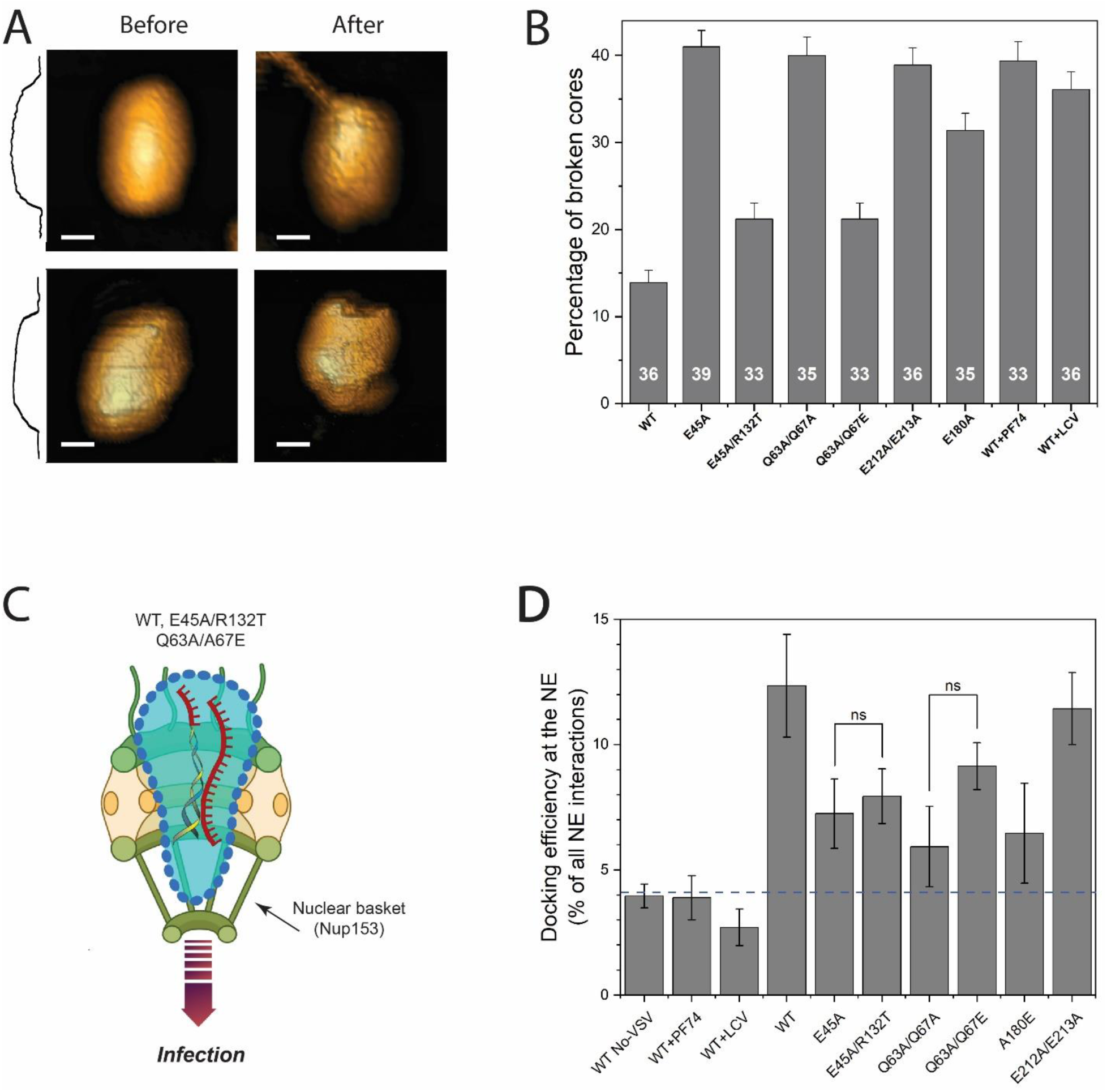
The effect of exposure to a high loading force on the structural integrity of cores. (A) Topographic AFM images of two representative WT cores before and after compression. Images were acquired using the QI mode operated at a 300 pN loading force. Typical cone-shaped cores are observed. A cross-section height profile along the length of the core is displayed. After compression, most cores (∼86%) remained intact. However, those that broke exhibited an opening at the narrow end of the core, distal from the compressed region. Scale bars are 50 nm. (B) The percentage of broken wild-type (WT) and mutant cores. Cores were examined in the presence of IP6 (100 μM). WT cores were also treated with the capsid-inhibiting antiviral drugs PF74 (1.25 μM) or Lenacapavir (LCV; 500 pM). The number of cores analyzed is indicated inside each bar. Core breakage was determined by AFM imaging of the cores after they were compressed by a loading force of 5 nN. Error bars represent standard error of the mean. (C) Model depicting the requirement for core elasticity in HIV-1 nuclear entry. Partially reverse transcribed cores interact with the external face of the NPC and undergo reversible deformation to insert into the pore. Capsid interactions with Nup153 promote passage through the pore. (D) Quantification of WT and mutant HIV-1 docking at the nuclear envelope (NE) in TZM-bl cells. The background level of NE interactions is shown as a blue dashed line, determined from endosome trapped bald-HIV-1 (no-VSV) infections that cannot engage NPCs. Data represent mean values acquired from 10 nuclei, and >600 tracks for each condition. Statistical significance was determined using non-parametric Mann-Whitney Rank Sum Test. Error bars represent standard error of the mean.

To investigate the potential role of elasticity in HIV-1 infection, we measured the structural recovery of cores from CA mutants that exhibit impaired nuclear entry and quantified the percentage of broken cores following compression as a measure of inelasticity/brittleness (Fig. 5B). Representative AFM images of intact and broken cores following compression are shown in Extended Data Fig. 9. In contrast to the wild type, E45A mutant cores were more brittle (41% of cores were broken following compression). By contrast, E45A/R132T cores were less brittle, exhibiting breakage (21%) similar to wild type cores, indicating that the addition of the suppressor mutation restored elasticity to the E45A mutant capsid narrow end. Cores from the nuclear entry-impaired mutants E212A/E213A, Q63A/Q67A, and E180A mutant cores were also brittle, exhibiting 40%, 39%, and 32% breakage upon compression, respectively. Substitution of glutamic acid for alanine at position 67 (Q63A/Q67E) restored elasticity to the wild type level (21% of broken cores). Moreover, addition of PF74 or LCV to wild type cores rendered them brittle as reflected by increased breakage upon compression (to 39% and 36% for PF74 and LCV, respectively). Thus, capsid-targeting small molecules reduce HIV-1 core elasticity at concentrations that inhibit nuclear entry. As previously shown for E45A and Q63A/Q67A ^30^, infection by the nuclear entry-impaired mutants E180A and E212A/E213A was resistant to the inhibitors. Like E45A/R132T, the pseudorevertant Q63A/Q67E was sensitive (Extended Data Fig. 10).

Our finding that the CA substitutions reduced HIV-1 core elasticity provides a compelling mechanistic explanation for the nuclear entry defect associated with the E45A and Q63A/Q67A mutant virus and their suppression by R132T and Q67E. By restoring elasticity, the R132T or Q67E mutations may permit penetration of the nuclear pore complex to enable interaction of the capsid with the nuclear basket protein Nup153, thus promoting nuclear translocation (Fig. 5C). This model is consistent with the known PF74 resistance and susceptibility of the E45A and E45A/R132T mutants, respectively, as well as the effects of these mutations on HIV-1 infection of nondividing cells ^29,39^. It may also account for the reduced effect of Nup153 depletion on infection by the E45A, Q63A/Q67A, E180A and E212A/E213A mutants ^1^ (Fig. 1E).

Mutants with brittle cores may also exhibit impaired nuclear entry owing to premature cytoplasmic breakage events. To address this issue, we employed live-cell fluorescence imaging microscopy to assess the docking efficiency of these cores at the nuclear lamina (Fig. 5D). Our findings revealed that all mutant cores successfully reach the nucleus, surpassing the threshold for random interactions. Notably, the docking efficiency of the brittle E45A and Q63A/Q67A mutants closely mirrored that of their elastic counterparts, E45A/R132T and Q63A/Q67E, respectively. Furthermore, the docking efficiency of brittle E212A/E213A cores was similar to that of wild-type (WT) cores. Collectively, our results establish that core elasticity does not correlate with the changes in docking efficiency. Additionally, our findings emphasize that, although all core mutants successfully reach the nuclear envelope, the key disparity lies in their ability to penetrate the nuclear pore, a characteristic exclusively exhibited by elastic cores. Nonetheless, we cannot exclude the possibility that the brittleness of the mutant cores leads to premature breakage of mutant capsids during reverse transcription in the cytoplasm, which may impair docking at the NPC and nuclear entry and result in activation of the cytoplasmic DNA sensor cGAS ^40^.

Our observation that addition of PF74 and LCV decreases the elasticity of the wild type capsid further suggests that elasticity is critical for HIV-1 nuclear entry. CA mutations that decrease the elasticity of the viral core confer resistance to these two inhibitors, likely by rendering the virus unable to utilize the nuclear pore complex. The elasticity-reducing mutations we have identified reside at the three major intersubunit interfaces in the capsid lattice, indicating that each interface is critical for elasticity. The suppressor mutations restored capsid elasticity, infectivity, nuclear entry, and PF74 and LCV sensitivity. We suggest that the HIV-1 capsid inhibitors PF74 and LCV inhibit nuclear entry by decreasing the elasticity of the viral capsid in addition to impairing its interactions with nucleoporins.

In conclusion, HIV-1 cores are highly elastic, endowing the core with the ability to undergo reversible deformation without structural failure. To our knowledge, this is the first reported example of such a highly elastic viral capsid. Recent structural studies of NPCs within cells have revealed that the diameter of the nuclear pore is sufficient to permit passage of the intact core ^4^. Nonetheless, the pore is normally filled with a high density of FG domains of nucleoporins. Also, NPCs in cells are dynamic, undergoing dilation and constriction ^41^. Docking of the reverse-transcribing viral core to the NPC may signal both the core and the NPC to dynamically adapt to one another, thus promoting passage of the core through the pore.

## Materials and Methods

### HIV-1 core production and core purification

Human embryonic kidney (HEK) 293T cells were used for the preparation of HIV-1 pseudovirus particles as previously described ^38^. Briefly, ∼10^6^ HEK 293T cells were cultured in a 15 cm dishes and transfected with 2.5 µg of Env-defective variants of the full-length HIV-1 construct R9 using 10 µg of polyethyleneimine (PEI; branched; MW ∼25000; Sigma-Aldrich). After 20 h, the cell medium (Dulbecco’s modified Eagle medium, supplemented with 10% heat-inactivated fetal bovine serum, 1% penicillin-streptomycin, and 1% L-glutamine) was replaced with fresh medium. After 6–7 h, the virus-containing supernatant was harvested, centrifuged at 1,000 rpm for 10 min, and filtered through a 0.45 µm pore size filter. The supernatant was concentrated using ultracentrifugation in an SW-28 rotor (25,000 rpm, 2 h at 4°C) on a 60% OptiPrep density cushion (Sigma-Aldrich). The pelleted viruses were resuspended in 10 mL TNE buffer (50 mM Tris-HCl, 100 mM NaCl, 0.1 mM EDTA; pH 7.4), and added to 100-kDa molecular mass cutoff Vivaspin 20 centrifugal concentrators (100,000 MWCO; Sartorius AG, Germany). The mixture was centrifuged twice at 2,500 g for 25–30 min at 4°C until the supernatant level in the concentrator reached a final volume of 300–350 µL.

Virus cores were isolated from the concentrated virus-containing supernatant using a previously described protocol ^38^. Purified HIV-1 virus particles were mixed with 1% Triton-X 100 diluted in 100 mM of 3-(N-morpholino) propane sulfonic acid (MOPS) buffer (pH 7.0) and incubated on ice for 2 min. The mixture was centrifuged at 13,800 g for 8 min at 4°C. After removing the supernatant, the pellet was washed twice using ∼80 µL of MOPS buffer and repelleted. The final pellet was resuspended in a 10 µL MOPS buffer. A fresh batch of cores was prepared on the same day as each of the AFM measurements. When specified, IP6, PF74, or Lenacapavir were added to the buffer at a final concentration of 100 µM, 1.25 µM, or 500 pM, respectively.

### AFM measurements and analysis

Samples for AFM measurements and analysis were prepared as previously described ^38^. Briefly, 10 µL of isolated HIV-1 cores were incubated for 45 min at room temperature on hexamethyldisilazane (HMDS)-coated microscope glass slides (Sigma-Aldrich) in a mildly humid chamber. AFM measurements were performed on the adhered sample without fixation. Each experiment was repeated at least three times, each time with independently purified pseudoviruses. IP6 was received as a gift from the Leo laboratory. PF-3450074 was synthesized by the Vanderbilt Institute of Chemical Biology Synthesis Core, Vanderbilt University, Nashville, TN. Lenacapavir (GS-6207, Gilead) was purchased from MedChemExpress (MCE), USA. All measurements were carried out with a JPK Nanowizard Ultra-Speed atomic force microscope (JPK Instruments, Berlin, Germany) mounted on an inverted optical microscope (Axio Observer; Carl Zeiss, Heidelberg, Germany). Silicon nitride probes (mean cantilever spring constant of 0.12 N/m; DNP, Bruker, Germany) were used. Topographic images were acquired using the quantitative imaging (QI) mode at a rate of 0.5 lines/s and a loading force of 300 pN and rendered using the WSxM software (Nanotec Electronica).

### Stiffness measurements

Core stiffness was obtained by operating the AFM in nanoindentation mode as previously described ^38,42^. Briefly, to determine the stiffness of each core, 20 force-distance (FD) curves were obtained from 24 different points on the core surface for a total of 480 FD-curves per core. To confirm that the capsid remained stable during the entire indentation experiment, each experiment was analyzed by the plotting individual point stiffness as a histogram and as a function of the measurement count. Samples whose point stiffness values decreased consistently during measurement, indicative of irreversible deformation, were discarded. The number of discarded FD-curves was 1 from each sample (corresponding to ∼3.5%). Importantly, this small number of discarded cores was similar for all samples, regardless of their elasticity. The maximal indentation of the sample surface was 4 nm which corresponds to a maximum loading force of 0.2–1.5 nN. Each FD-curve was shifted to set the deflection in the non-contact section to zero (baseline). The set of FD-curves was then averaged. The measured stiffness was mathematically derived from the slope of the FD-curves. Core stiffness was computed using Hooke’s law, assuming that the experimental system can be modeled as two springs (the core and the cantilever) arranged in series. The spring constant of the cantilever was determined during experiments by measuring the thermal fluctuations. The data were analyzed using JPK Data Processing and MATLAB (The Math Works, Natick, MA) software. To test the mechanical strength of the core, a single FD curve was acquired from a single central point on the surface of the core. The maximum loading force was set to either 5 nN, 10 nN or 20 nN.

### Volume recovery measurements

The elasticity of the core was assessed in terms of volume recovery after forcibly compressing the structure. The entire core was imaged using the AFM operated in the QI mode using a low maximal force of 300 pN. A small rectangular region on the main body of the core was then selected and scanned. The chosen region was then compressed by re-scanning it at a maximal loading force of 5 nN. Higher maximal loading forces frequently resulted in the detachment of cores from the glass substrate as evident by the disappearance of the core in the consequent images. The force was then lowered back to 300 pN, and the region was repetitively imaged for a total duration of 5–6 minutes. At the end of the experiment, the entire core was imaged at a 300 pN loading force. The collected set of images was then processed using an in-house MatLab script to calculate the volume underneath the selected region. The initial volume prior to compression was set to 100%, and the rest of the calculated volumes were normalized accordingly. The obtained volume values were then plotted against time. The initial and final images of the entire core were used to determine if the core underwent structural breakage upon compression.

### HIV-1 infection assays

Infectivity was quantified by titration of GFP reporter viruses bearing the indicated amino acid substitutions in CA. Virus stocks were prepared by cotransfection of 3 μg of Env-defective HIV-GFP proviral plasmid ^43^ with 0.5 μg pHCMV-G ^44^, generating VSV-pseudotyped particles. After culturing for 40-48 h, virus-containing culture supernatants were collected, clarified by passing through 0.45 μm syringe filters, frozen in aliquots, and stored at -80°C. Virus concentrations were determined by p24 ELISA ^45^. Infections were performed by inoculating 20,000 Hela cells seeded one day previously in 48-well plates with 0.25 ml of virus dilutions prepared in complete medium containing 20 μg/ml DEAE-dextran. Cultured were incubated for two days and cells detached by treatment with trypsin, collected, fixed overnight in 0.4% paraformaldehyde, and analyzed for GFP expression by flow cytometry. Infectivity was determined as % GFP^+^ cells in the culture normalized by the concentration of p24 in the inoculum with wells exhibiting between 1% and 20% GFP^+^ cells. To assay inhibition of infection by PF-3450074 (PF74) and Lenacapavir (both purchased from MedChemExpress), the compounds were added to a dilution of each virus that gave 5-10% infected cells in the absence of inhibitor. Two days later, the cells were fixed and analyzed for infection by flow cytometry.

In experiments involving cell cycle arrest, Hela cells (40,000) were seeded in medium containing aphidicolin (1 μg/ml). The next day, medium was removed, and aphidicolin was included in the virus inocula. The following day, the medium was replaced with fresh medium lacking aphidicolin. In experiments involving Nup153 depletion, cells were first transfected with control or Nup153-targeting siRNA duplexes as previously described ^32^. Two days later, cells were replated for infection assays and for immunoblotting to confirm the knockdown efficiency.

To isolate the Q63A/Q67E pseudorevertant, cultures of MT-4 cells (200,000) in 1 ml of medium were inoculated with R9.Q63A/Q67A mutant virus (25 ng p24) encoding a Vpu truncation at amino acid 35. The cultures were maintained and assayed for HIV-1 accumulation by p24 ELISA every four days. After approximately two weeks, p24 was detected in the mutant virus culture. DNA was extracted and purified using a DNAeasy kit (Qiagen), and the product used as a template for PCR amplification of the Gag coding region. PCR products were gel purified, sequenced, and cloned into the wild type R9 and HIV-GFP reporter virus plasmids. The PCR-amplified regions of the plasmids were sequenced to confirm identity.

For quantifying reverse transcription and nuclear entry, monolayers of 200,000 Hela cells in 12-well dishes were inoculated with 0.5 ml of virus, corresponding to 20 ng of p24, that had been previously treated with DNaseI to reduce the level of contaminating plasmid DNA. Infection was performed in medium containing 20 μg/ml DEAE-dextran. When required, Efavirenz or Raltegravir was added to diluted viruses to a final concentration of 1 μM. Cultures were incubated for 16 hours, cells were collected, and DNA was purified on silica columns ^46^. DNA was eluted in 100 μl of water, quantified by spectrophotometry, and assayed for late reverse transcripts and 2-LTR circles by quantitative PCR with standard curves generated from dilutions of appropriate plasmids.

### Simulated AFM nanoindentation experiments

#### HIV-1 CA conical capsid construction

The coarse grained HIV-1 conical capsid was modeled using a methodology we developed previously^47^. Details of the coarse graining methodology, as well as parameterization and model validation, were previously reported ^47^. The system includes models of sodium, chloride, and host-factor inositol hexakisphosphate (IP6), yielding 150 mM NaCl ionic strength and 253 IP6 molecules bound to central pores of capsomers ^48^. The charge-neutral, ionized and IP6-bound capsid was then equilibrated for 500 ns using NAMD3 ^49^.

#### Molecular dynamics simulations

All molecular dynamics (MD) simulations were performed with the nanoscale molecular dynamics (NAMD) engine ^49^. NAMD 2.14 was utilized for energy minimization and the fully GPU-resident NAMD 3 was utilized for NVT equilibration and constant velocity steered MD (cv-SMD) simulations. Simulations of wild type capsids utilized a time step of 48 femtoseconds, whereas mutant E45A and E45A/R132T capsid simulations utilized a time step of 40 femtoseconds. Long range electrostatic evaluation was accomplished via particle mesh Ewald (PME) ^49,50^. The non-bonded potential excluded bonded neighbors (1-2 exclusion policy) and utilized a cutoff of 2 nm. Short range non-bonded interactions were computed at every time step, while long range interactions were computed every other time step. The discrete grid used in PME for interpolating charges was given a spacing of 2 Å, employing 8^th^ order interpolation. Temperature was enforced via Langevin dynamics, with a target temperature of 298 K. All beads besides the AFM tip were coupled to the thermal bath with a γ coefficient of 2 ps−1. A dielectric constant of 80 was employed to mimic charge screening by solvent molecules ^51^.

#### In silico AFM simulations

For AFM simulations, constant velocity pulling was applied to a dummy atom with a velocity of 1.5 × 10^−6^ Å per time step, or 3.125 nm per microsecond. A harmonic force constant of 12 kcal/mol was employed to restrain the 13 nm AFM tip’s center of mass to the dummy atom. Additionally, a transverse harmonic force constant of 1 kcal/mol was employed. Data from the AFM simulations, namely forces acting on the AFM tip throughout simulation as well as the tip’s position data, were output with a frequency of 48 picoseconds, or every 100 time steps (A detailed description of the AFM simulations is provided in the Extended Data).

### Live-cell imaging of HIV-1 docking at the nuclear envelope and nuclear import

Single HIV-1 infection in live cells was visualized, as previously described ^13,52^. In brief, 8·10^4^ or 5·10^5^ TZM-bl cells that stably express the emiRFP670-laminB1 nuclear envelope marker was respectively plated in a 8-well chambered slide (#C8-1.5H-N, CellVis) for single time-point nuclear import experiments, or in a 35mm 1.5 glass bottom dish (#D35-20-1.5-N, CellVis) for NE-interactions experiments. Aphidicolin (Sigma Aldrich) 10 µM was added to cells to block cell-division. About 14h later the cells were infected (MOI 0.2, for NE interactions (or) MOI 1 for nuclear import) with indicated WT and mutant HIVeGFP particles fluorescently tagged with Vpr-integrase-mNeongreen (INmNG) to label the viral core. Virus binding to cells was augmented by spinoculation (1500×g for 30 min, 16°C), and virus entry was synchronously initiated by adding pre-warmed complete DMEM medium containing Aphidicolin (10 µM) to samples mounted on a temperature- and CO_2_-controlled microscope stage. 3D time-lapse live cell imaging was carried out on a Leica SP8 LSCM, using a C-Apo 63x/1.4NA oil-immersion objective. Tile-scanning was employed to image multiple (4×4) fields of view. Live-cell imaging was performed starting from 0.5 – 16 hpi by acquiring 9-11 Z-stacks spaced by 0.8 μm every 2.5 min. The Adaptive Focus control (AFC, Leica) was utilized to correct for axial drift. 512 x 512 images were collected at 180 nm pixel sizes at a scan speed of 1.54 µs pixel dwell time. Highly attenuated 488 and 633 nm laser lines was used to excite the INmNG and emiRFP670-Lamin B1 fluorescent markers and their respective emission was collected between 502-560 nm and 645-700 nm using GaSP-HyD detectors. For assessing HIV-1 nuclear import in a fixed time-point (8 hpi), more stringent imaging conditions were used i.e., 2x laser power, with 0.12 μm/pixel sizes and 0.5 μm spaced z-stacks. 3D-image series were processed off-line using ICY image analysis software (http://icy.bioimageanalysis.org/) ^53^.

### Image analyses

HIV-1 nuclear import in fixed time-point 3D z-stack images were analyzed using an in-house script in the ICY protocols’ module as described in ^52^. For HIV-1 docking at the NE analysis, we selected >10 cells from each independent live-cell experiments that showed minimal lateral movements. Additionally, the movies were cropped, and analysis was performed only for the central 8h (between 4-12hpi) to overcome caveats associated with initial particle attrition and latter cellular motions. The central z-planes (2x planes, 1.6 um) of these nuclei was extracted and projected to a 2D-stack, which was additionally drift corrected using Fast4Dreg in Fiji ^54^. Images were denoised using gaussian filters, and the laminB1 fluorescence signals was used to mask the pixels occupied by the NE as a region of interest. The HIV-1 INmNG puncta detected within this NE mask was tracked in an automated fashion in ICY bio-image analysis single particle tracking suite, and each individual trajectory was analyzed manually for NE-interaction and docking behavior. Segments within individual HIV-1 trajectories that coincide with the NE mask for >2 frames (>5 min) were considered as NE-interactions, and the segments of tracks when a single HIV-1 particle remained within a 2-pixel (360 nm) radius for 3 or more frames (>7.5 min) was considered as HIV-1 core docking at the NE. The particle displacement along the NE was determined by a rolling average of particle displacement. The 360nm radius was used taking into consideration the lateral diffraction limited confocal resolution and the pixel sizes in our time-lapse movies, which is 250nm and 180nm, respectively. These stable docking events, as a fraction of total NE-interactions (total number of HIV-1 tracks, including track segments that was not localized within the 360nm window) obtained in each individual nuclei over the 8h time-window was analyzed and plotted. As controls, the interactions between bald endosome trapped HIV-1 (no-VSV) infections were analyzed and the background in detecting non-specific overlaps as HIV/NE-docking was estimated to be 4.2%.

## Author Contribution

A.D., A.B., J.R.A.M, J.S., J.H., C.T., and C.A. performed research and analyzed data, A.D., A.B., S.H., J.R.A.M., and A.C.F analyzed data, C.A, J.P., A.C.F., and I.R. designed research and wrote the paper.

## Supporting information

Supplemental figures, discussion and methods

## Acknowledgments

We thank Harrison Bracey and Ashley Fulmer for technical assistance and Alan Engelman for helpful comments. This work was supported in part by the Israel Science Foundation grant 418/21 (Rousso Lab), by NIH R01-AI157843 and U54-AI170791 (Aiken and Perilla Labs), NIH U54 AI170855 and R01-AI148382 (Francis lab). C.T. received support from NIH 5R25AI164610 and a Patrick and Eloise E. Wieland Fellowship. This work used Delta at NCSA and Expanse at SDSC through allocation MCB-170096 from the Advanced Cyberinfrastructure Coordination Ecosystem: Services & Support (ACCESS) program, which is supported by National Science Foundation awards #2138259, #2138286, #2138307, #2137603, and #2138296.

## Data availability

The data that support the findings of this study are available in the manuscript and its Supplementary Information and from the corresponding authors on reasonable request.

