## Supplemental figures, discussion and methods for "Elasticity of the HIV-1 Core Facilitates Nuclear Entry and Infection"

[1]Ben-Gurion University of the Negev, Department of Physiology and Cell Biology, Beer Sheva, Israel; [2]University of Delaware, Department of Chemistry and Biochemistry, Newark DE, USA; [3]Florida State University, Institute of Molecular Biophysics, Tallahassee, FL, USA; [4]Vanderbilt University Medical Center, Department of Pathology, Microbiology and Immunology, Nashville, TN, USA; [5]Florida State University, Department of Biological Sciences, Tallahassee, FL, USA

| | Pages |
|---|---|
| Supplemental Figures | 2-12 |
| Supplemental MD discussion | 13-15 |
| Supplemental Methods | 16-19 |

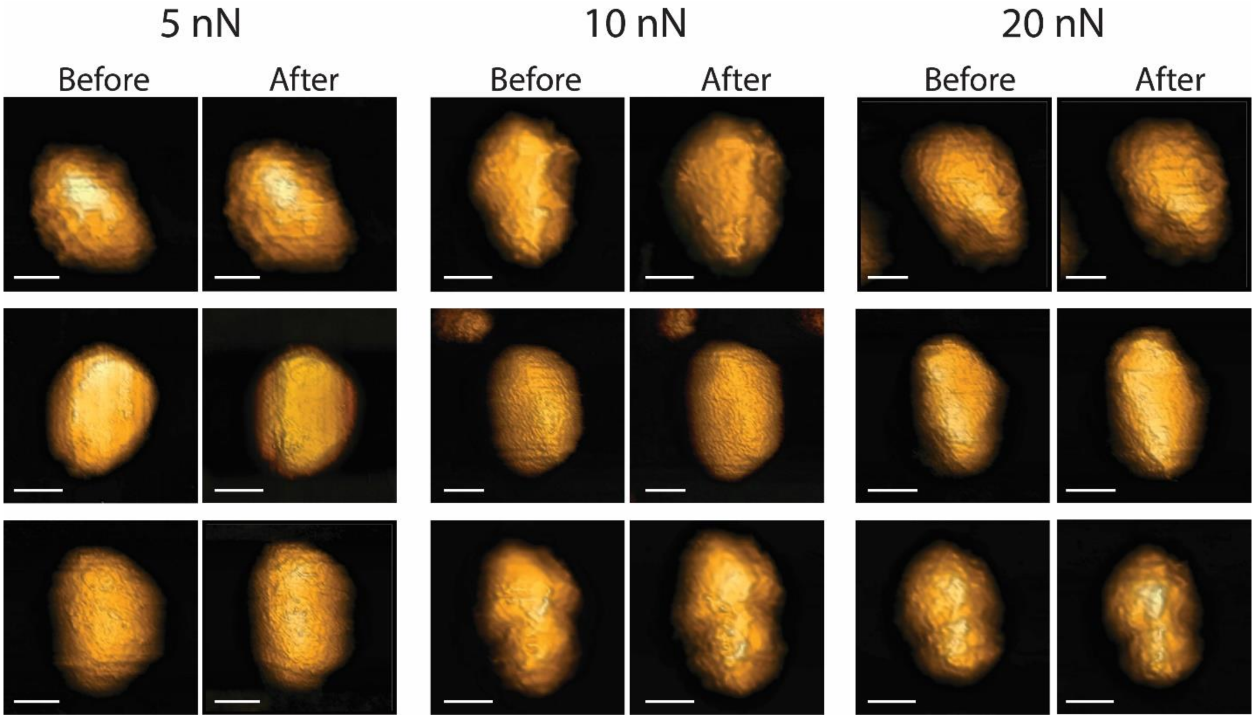


Extended Data Fig. 1: Topographic AFM images of isolated IP6-treated WT cores before and after FD curves applied at maximal loading force of 5 nN, 10 nN or 20 nN. Three representative pairs of images for each maximal loading force. All images were acquired using the QI mode at a maximal loading force of 300 pN. Scale bars are 60 nm.

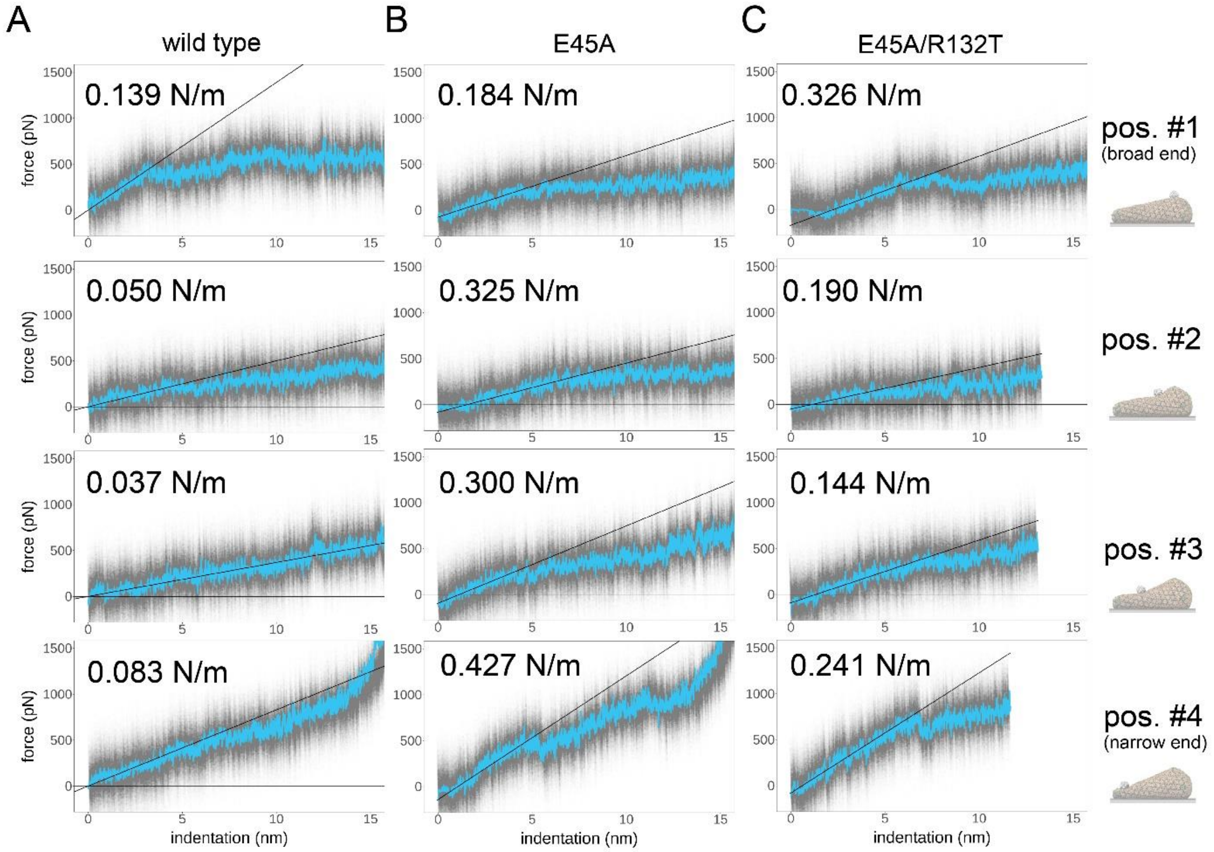


Extended Data Fig. 2 – Simulated atomic force microscopy indentations (3.125 nm/microsecond probe velocity) for **A** wild type, **B** E45A, and **C** E45A/R132T capsids. Each row of the composition represents a different probe location corresponding to those shown in Fig. 2 **B**-**E** and are labeled accordingly. For each plot, probe approach and initial adhesion to the sample surface were omitted, such that the x-axis shows sample indentation in units of nanometers. The linear fits of the first four nm of indentation, from which stiffness values were computed, are shown with the relevant stiffness value annotated. E45A and compensatory E45A/R132T mutant capsids are considerably stiffer than wild type in all locations probed. Wild type stiffness: 0.077 +/- 0.045 N/m; E45A stiffness: 0.309 +/- 0.099 N/m; E45A/R132T stiffness: 0.225 +/- 0.078 N/m.

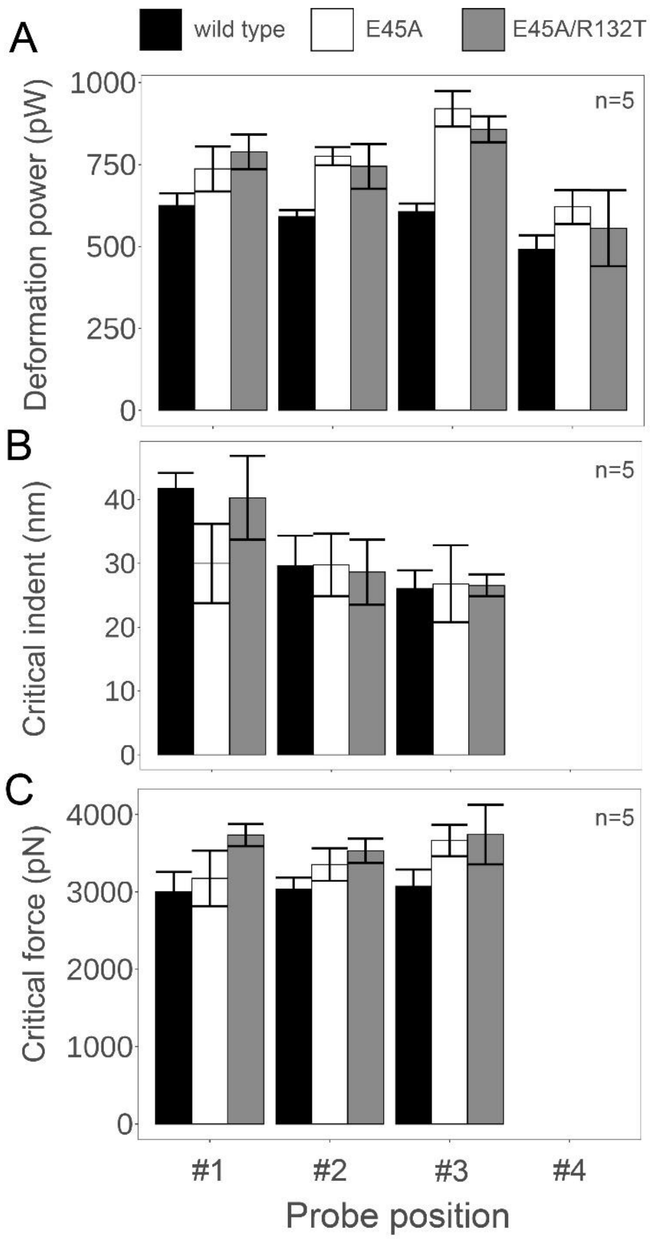


Extended Data Fig. 3 – Summary of high-velocity AFM simulation measurements to evaluate the material response of the HIV-1 capsid in the high-force regime. Shown are the values of (A) power, (B) indentation, and (C) force at which the capsid is ruptured. Mean values from n=5 simulations are shown, with error bars representing the standard deviations. For panels B and C, position #4 values are omitted; failure of the capsid lattice, and thus critical indentation and force values, is not well-defined due to the relatively small height of the sample in the narrow end.

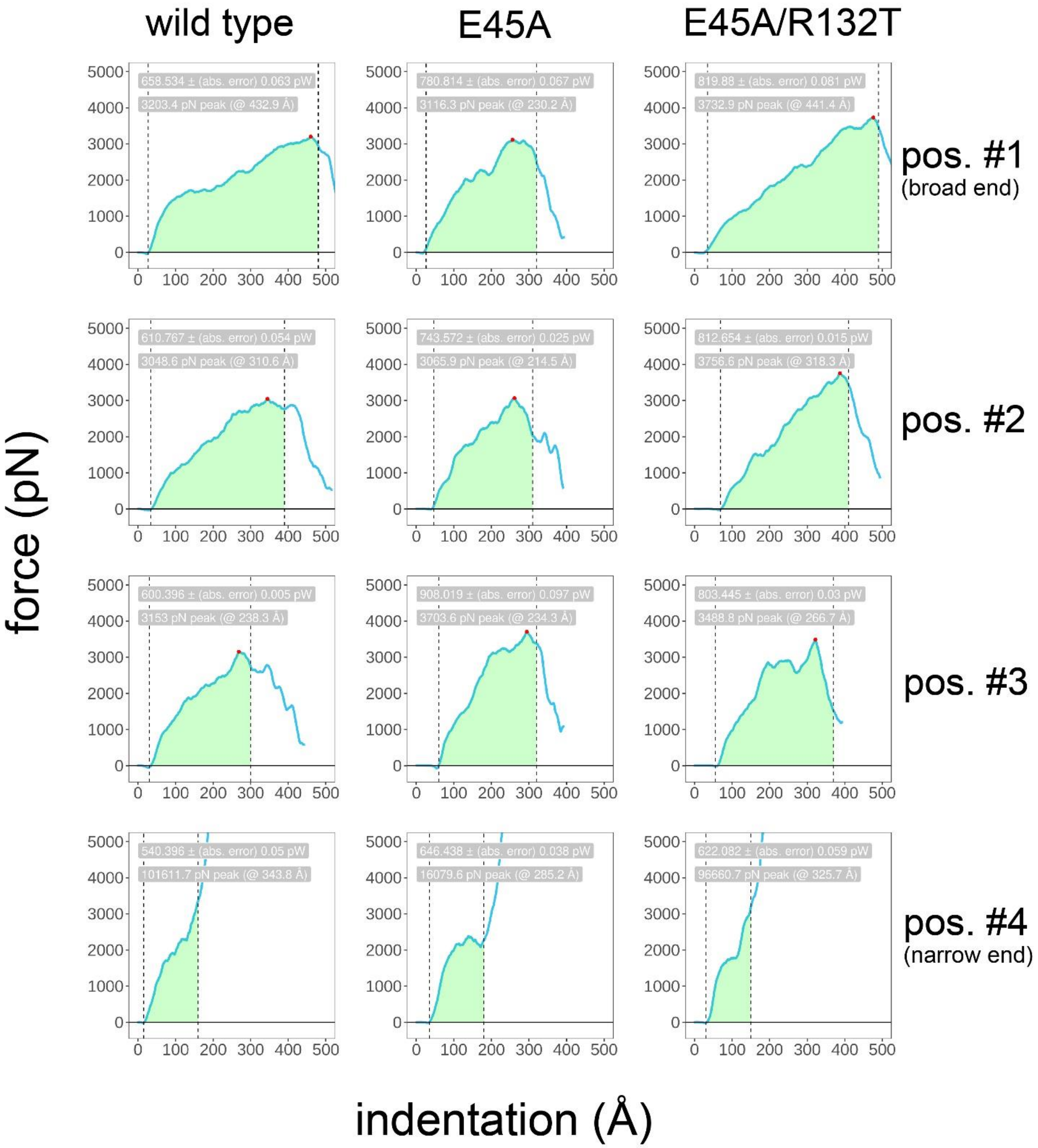


Extended Data Fig. 4 – One of five exemplary trial of high velocity (312.5 nm/microsecond) atomic force microscopy simulations for three simulated capsids: wild type, E45A, and E45A/R132T. Utilizing a higher probe velocity mimics high-force AFM experiments, where we observed deformation and failure of capsids conferring higher measured forces. In the case of simulations, these events were observed across smaller time intervals. Each column is labeled according to the relevant construct, and each row represents a different probe location along the length of the cone (corresponding to Figure 2**B-E**). Each plot is annotated with the power, determined by integration (shown in green), that is summarized in Figure S5. Integration bounds are shown, the lower of which was set at probe contact and the upper bound is set to encapsulate the yield (critical) force, annotated with a red dot. Power is considered as the time-normalized integral of each curve (green region), presented in units of picowatts with absolute integration error given. For the narrow end probe locations, position four, critical forces and indentation distances are ill-defined due to densification of the capsids. For the latter cases, we set the upper integration bound as immediately prior to densification. This enables the calculation of power but not critical forces or critical indentation values as shown in Figure S5 for the other three probe positions.

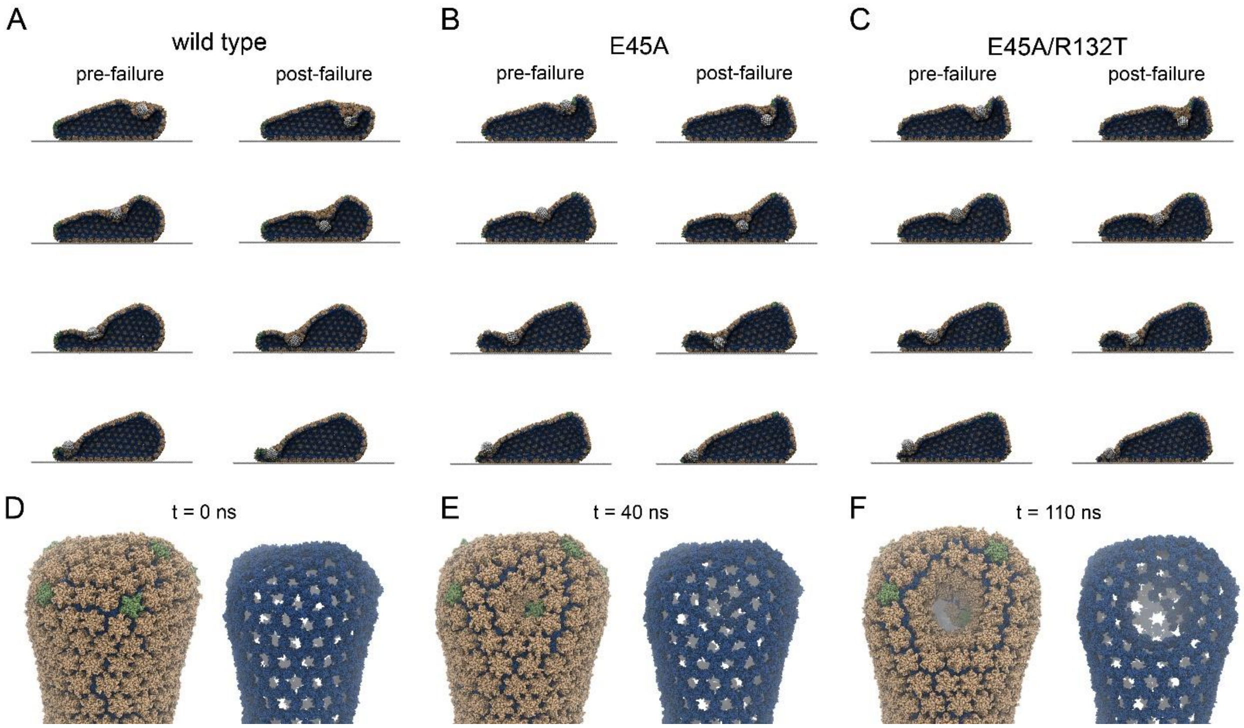


Extended Data Fig. 5 – Visualization of deformed and ruptured capsids from AFM simulations. Pre- and post-failure snapshots are shown for: (A) wild type, (B) E45A, and (C) E45A/R132T capsids, with each probe position shown. Panels **D**-**F** show a succession of close-up images of wild type capsid indentation and failure, including a complete view as well as a C-terminal domain only view, to highlight the separation of assembly interfaces. **E** shows indentation without significant separation of assembly interfaces, slight separation of trimer interfaces is visible. **F** shows the failure event fully manifest, where a large discontinuity is seen in both the complete and C-terminal domain views.

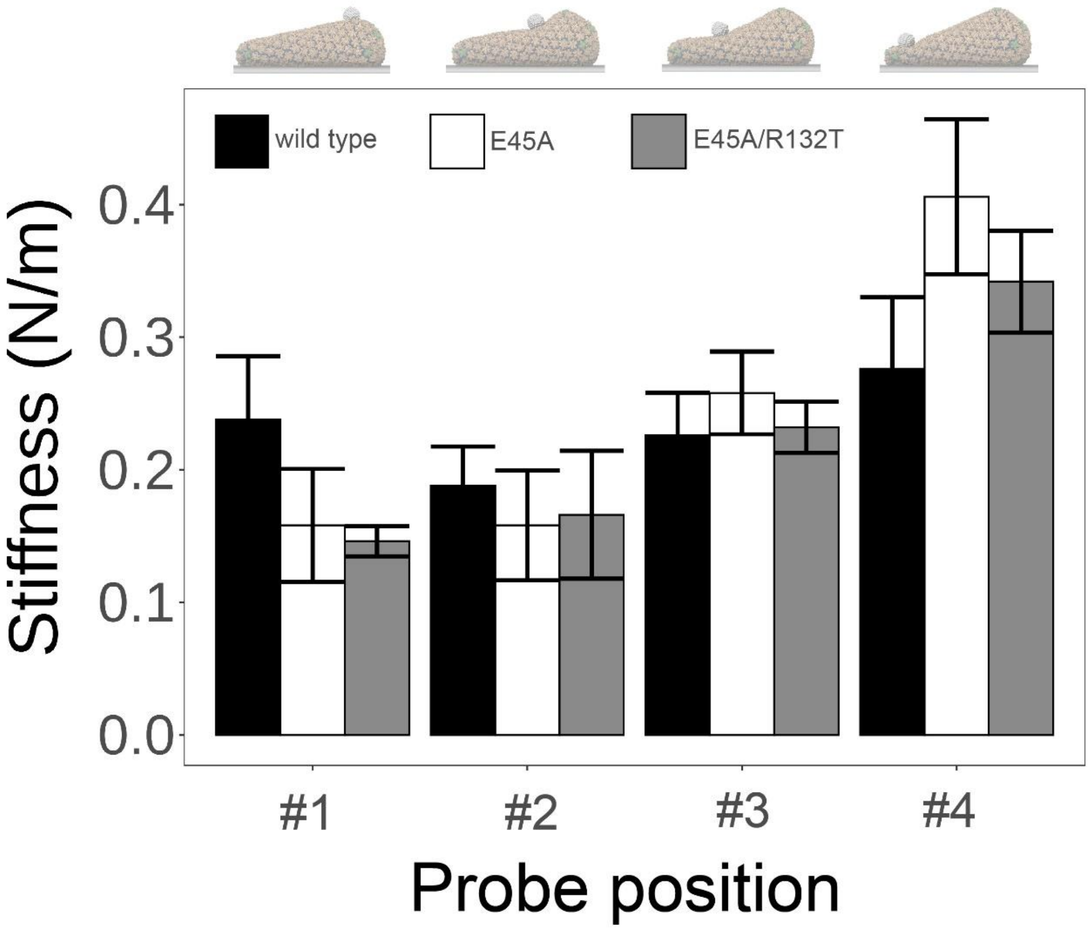


Extended Data Fig. 6 – Stiffness values from simulated AFM nanoindentation high velocity FD curves (as shown in Figure S5), where bars represent the mean of n=5 trials for each probe position, and for each construct: wild type, E45A and E45A/R132T, colored accordingly. Error bars represent the standard deviation of these five trials. Interestingly, using a significance threshold of 0.05, we see that wild type is significantly stiffer than either mutant at broad end position #1 (p-value = 0.02358 for E45A vs. wild type; p-value = 0.01089 for E45A/R132T vs. wild type). For narrow end position #4, E45A is significantly stiffer than wild type (p-value = 0.006611). For the remaining probe positions, all three constructs confer insignificant differences in stiffness, with p-values > 0.05. This conforms to experimentally derived stiffness trends of wild type, E45A and E45A/R132T capsids.

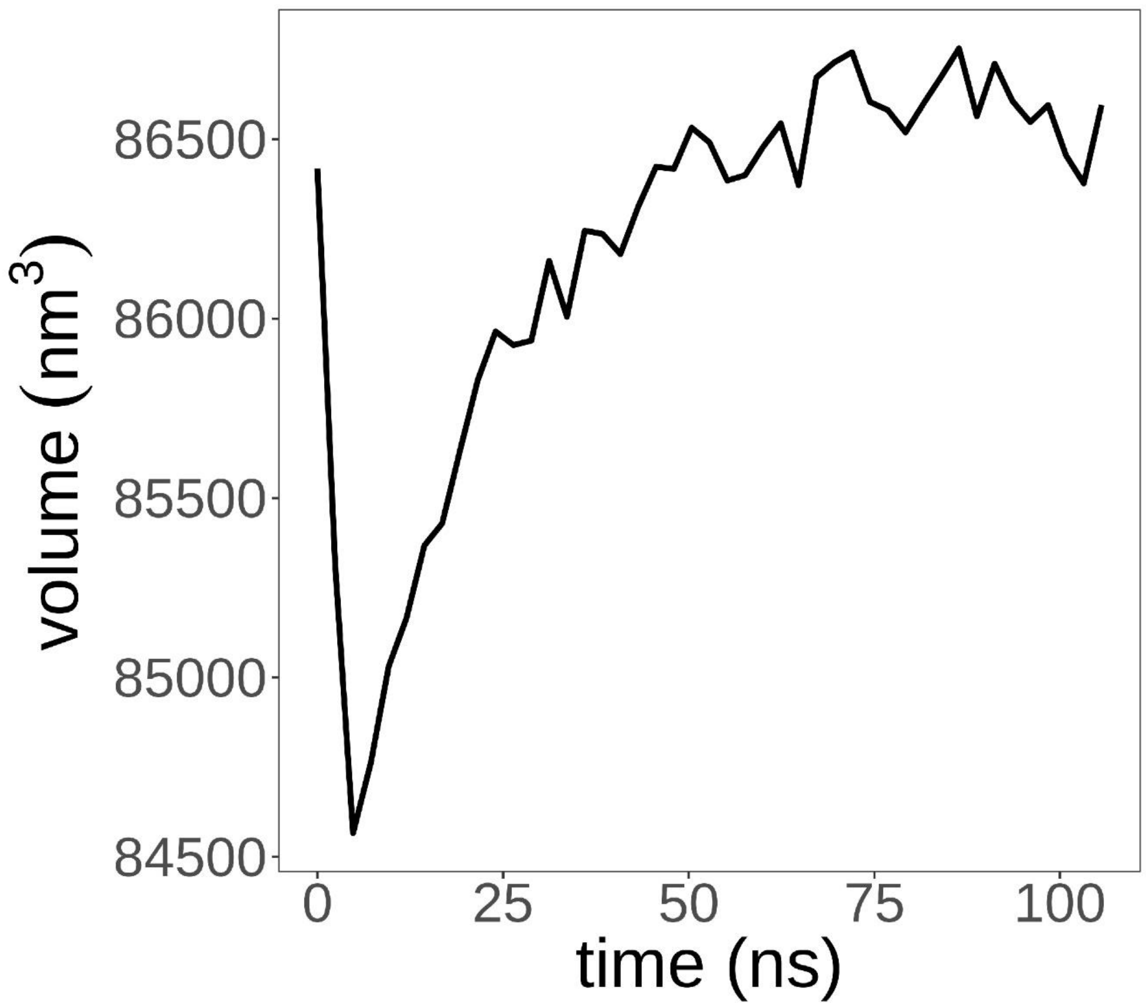


Extended Data Fig. 7 – Simulated AFM volume recovery experiment of a wild type capsid. Following a rapid indentation, the capsid is shown to recover its volume over a relatively short interval of 20 ns. This full and rapid recovery of capsid volume is consistent with experimental volume measurements shown in Figure 3C.

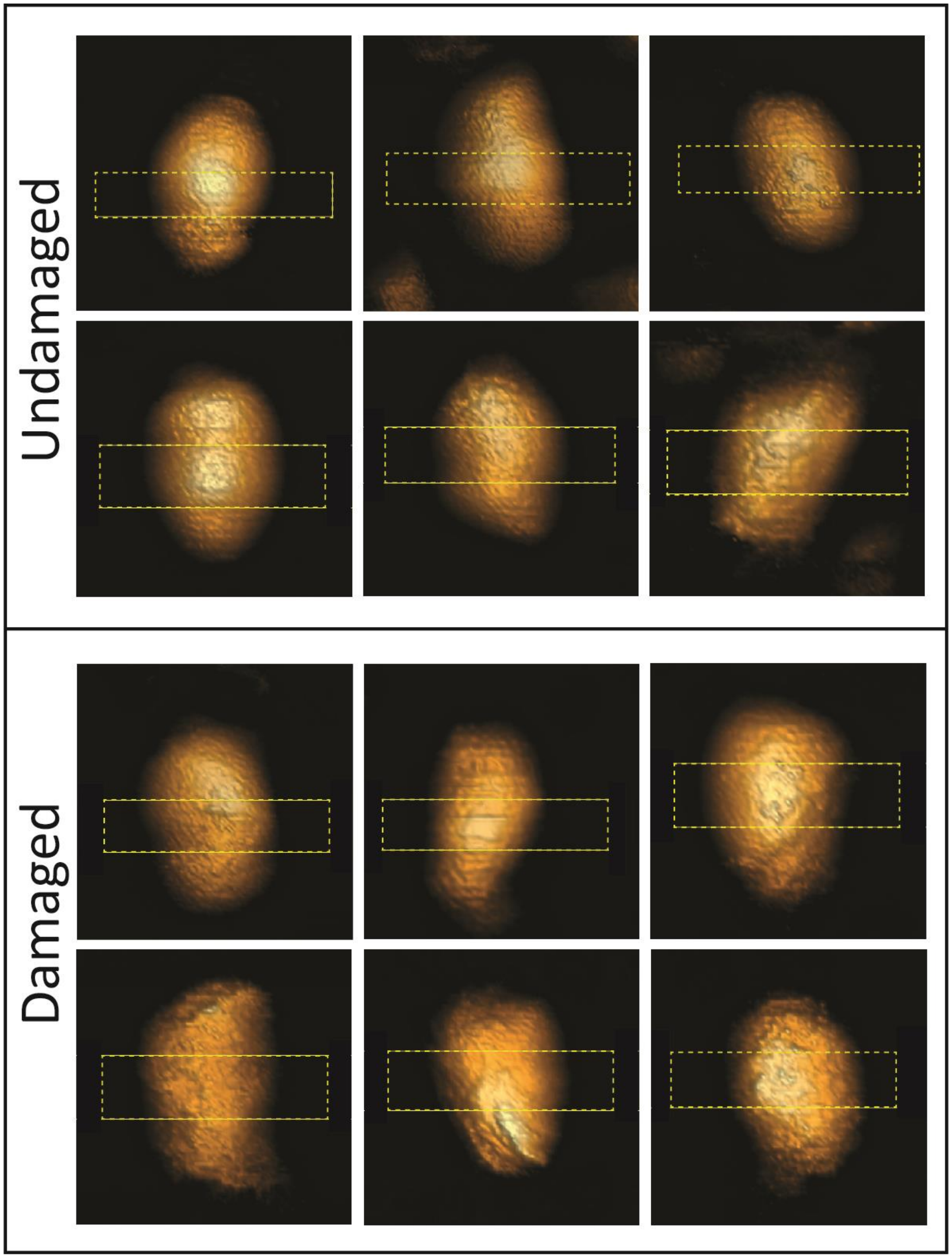


Extended Data Fig. 8: Topographic AFM images of isolated IP6-treated cores. All images were acquired using the QI mode at a maximal loading force of 300 pN. The region that was compressed at 5 nN loading force is labeled in dashed yellow rectangular.

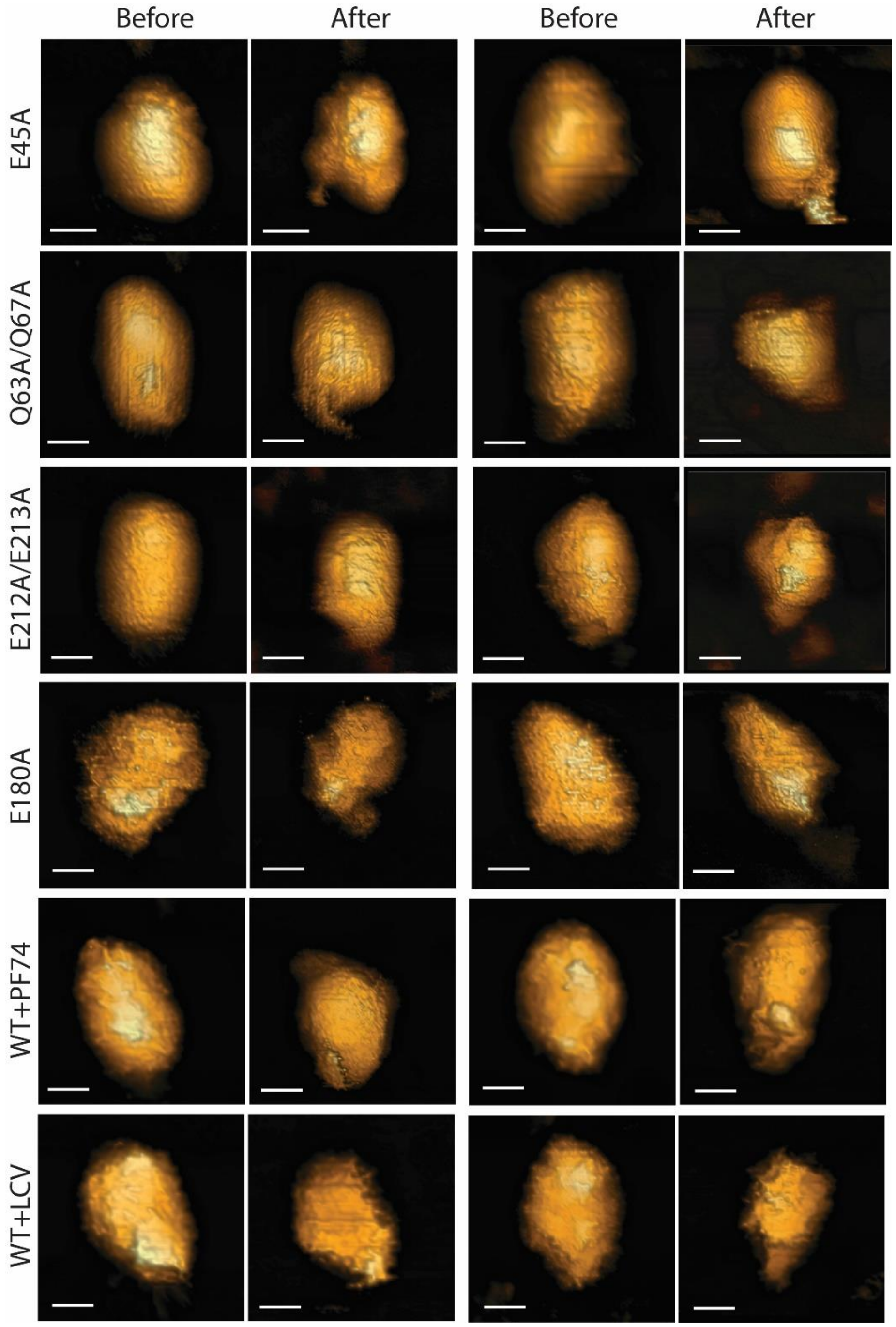


Extended Data Fig. 9: Topographic AFM images of broken IP6-treated cores before and following high-force (5 nN) compression. Three representative images for each mutant and WT treated with PF74 or LCV are shown. All images were acquired using the QI mode at a maximal loading force of 300 pN. Scale bars are 60 nm.

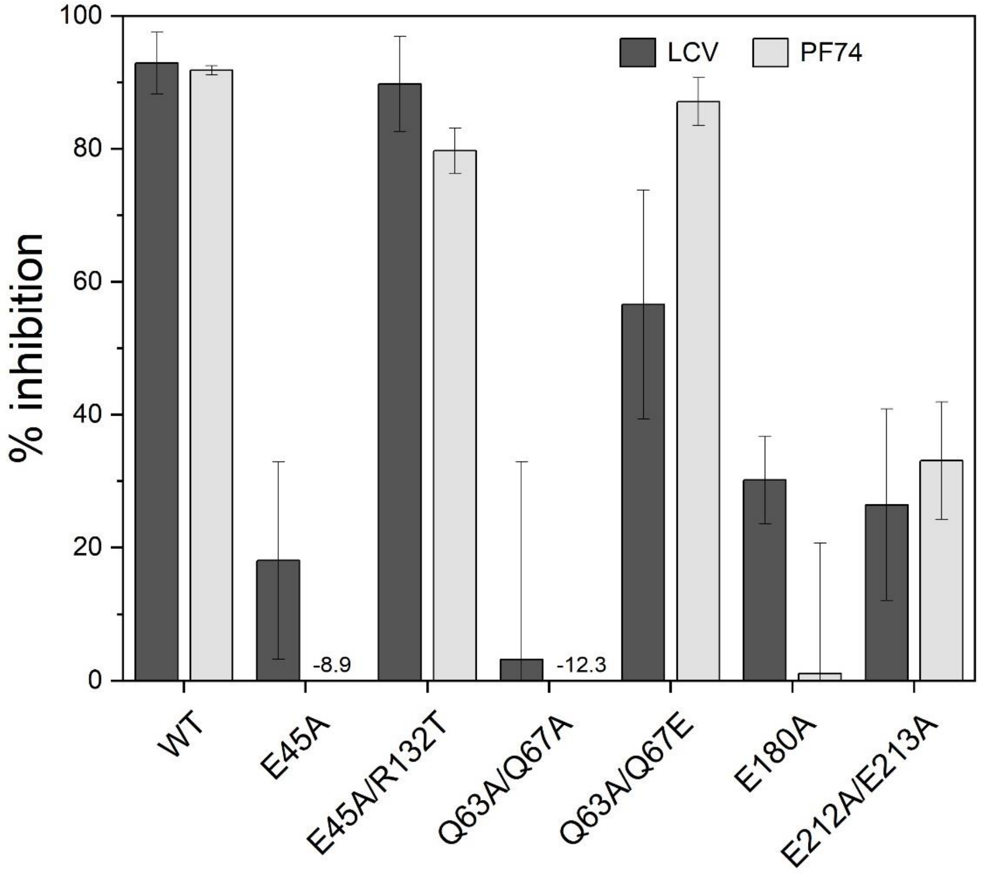


Extended Data Fig. 10: Effects of 0.25 nM Lenacapavir (LCV) and 1.25 μM PF74 on infection by HIV-1 CA mutants.

**Supplemental discussion of simulated AFM experiments**

Simulated AFM nanoindentation of the E45A and E45A/R132T mutant capsids revealed a striking increase in stiffness compared to wild type (Figure S4). Utilizing a 3.125 nm/μs probe velocity, the same as employed for wild type simulations (Figure 2B-E), we obtained significantly higher stiffness values for these mutants. For E45A: 0.309 +/- 0.099 N/m (n=4); for E45A/R132T: 0.225 +/- 0.078 N/m (n=4); and for wild type: 0.077 +/- 0.045 N/m (n=4). While our models revealed a decrease in stiffness of E45A/R132T vs. E45A, the E45A/R132T mutant core remains significantly stiffer than the wild type. The latter result is at odds with experimentally measured stiffness, where wild type is slightly stiffer than E45A and E45A/R132T mutants, and the mutant capsids are similar to one another.

We additionally subjected wild type and mutant capsid models to high-velocity (312.5 nm/μs) AFM simulations (Figure S5, S6). These simulations mimic high-force AFM experiments, where forces acting on the simulated AFM probes are higher, and we were able to record and observe failure events and thus determine critical forces and critical indentation values corresponding to failures. These simulations cover smaller timescales, enabling us to perform five repeated trials, amounting to 20 indentation simulations (Table S1) for each of the three simulated constructs, with n=5 trials per probe location. Strikingly, time-normalized integration of resulting force-displacement curves showed that E45A and E45A/R132T absorb considerably more power than wild type during indentation (Figure S5A). Moreover, broad end (position #1) deformation showed that E45A failed at significantly shorter indentation distances, whereas E45A/R132T sustained deformations similar to wild type (Figure S5B), despite also sustaining higher critical forces (Figure S5C).

We then computed mean stiffness across the five trials for each construct, treating each probe location independently rather than aggregating all probe positions into a single stiffness value (Figure S8). We found that the stiffness of each construct deviates significantly only at the highly curved broad and narrows ends (positions #1 and #4, respectively). Similar to the experimentally derived stiffness measurements showing wild type capsids are stiffer than E45A and E45A/R132T mutants, we see that the broad end (position #1) of the wild type capsid is the stiffest of the three constructs tested. Importantly, the flat mid-regions of the conical capsids yield similar stiffness measurements for all three constructs, conforming to experimentally derived trends in stiffness. While the exact distribution of probe locations employed for experimentally derived stiffness measurement is not known precisely, based on considerations of sample surface area we conclude that experimental nanoindentations disproportionately probe the broad end and mid-region of capsids (positions #1, #2 and #3), where wild type capsids confer a stiffer mechanical response and E45A and E45A/R132T mutants show similar responses.

| **Construct** | **Probe velocity (nm/µs)** | **Num. trials** |
|---|---|---|
| Wild type | 3.125 | 3 |
| E45A | 3.125 | 3 |
| E45A/R132T | 3.125 | 3 |
| Wild type | 312.5 | 5 |
| E45A | 312.5 | 5 |
| E45A/R132T | 312.5 | 5 |

Table S1 – Simulated AFM trials performed. For each construct, a trial is defined as a set of four probe positions, corresponding to Figure 2B-E.

**Supplementary methods**

*Molecular Dynamics*

*Preparation of AFM apparatus: baseplate and probe*

For the base plate, we utilize a rectangular, flat lattice of beads with van der Waals (vdW) radii of 1 nm. Bond distances of 1.5 nm are enforced with force constants of 10 $\mathrm{kcal \cdot mol^{-1}}$, lattice regularity and planarity are enforced via 90 and 180 degrees angle terms, respectively, with 100 $\mathrm{kcal \cdot mol^{-1}}$ force constants. The beads comprising the base plate are given a Lennard-Jones ε value of -1.5 $\mathrm{kcal \cdot mol^{-1}}$, to allow a weak and non-specific anchoring of the capsid to the plate. The AFM tip is modeled with inert beads of vdW radius 1 nm arranged in a cubic lattice, and by excluding all beads that are 6.5 nm away from the lattice's geometrical center. The resulting probe is a 13 nm diameter sphere where each bead comprising the sphere is bound to its neighbors with an equilibrium distance of 1.5 nm and a force constant of 10 $\mathrm{kcal \cdot mol^{-1}}$. Angle terms, either 90 degrees or 180 degrees, are enforced with force constants of 100 $\mathrm{kcal \cdot mol^{-1}}$. Four systems were built, where the tip is spaced equidistantly along the capsid's principal axis of inertia. Capsids were adsorbed for 150 ns prior to AFM simulations.

*Post-processing of force-profile curves*

For plotting, both the raw data and a windowed-average trace, employing a window size of 1,000 points, are shown (Figure 2). Linear fits for estimation of stiffness, in units of $\mathrm{N \cdot m^{-1}}$, were employed using the first 3 nm of indentation from each in silico AFM trajectory.

*Fluorescence Microscopy*

*Virus production*

Fluorescently tagged VSV-G pseudotyped HIV-1 particles was produced as described previously [1]. Briefly, envelope deleted pHIVeGFP CA wild-type (WT) and indicated CA mutants (MT) proviral backbone (2ug), VSV-G envelope (0.5μg) and Vpr-integrase fused to mNeonGreen (INmNG, 0.8μg) was mixed in jetPRIME buffer and 6ul of jetPRIME reagent, and transfected into HEK 293T cells plated at 80% confluency in a 6-well plate. Following a 6h incubation in a $CO_2$ incubator, the transfection medium was exchanged for fresh phenol-red minus DMEM complete with antibiotics and 10% FBS. Virus supernatants were collected after an additional 36h of incubation, clarified through a 0.45 μm filter and quantified for RT-activity, aliquoted and stored at -80°C until use.

*Tracking the interactions between HIV-1 capsid mutants and the nuclear pore complex*

In prior work, others including our group used fast-temporal imaging of HIV-1 capsid interactions with the nuclear envelope (NE) in living cells [1,2]. In these experiments, interaction of a mutant capsids K203A or E45A were negligibly detected, primarily due to the imaging experiment which spanned for only a short time-window (2h). Moreover, it was not feasible to estimate fraction of capsids that dock at the NE during the entry time course of infection. To address these caveats, and to unbiasedly characterize HIV-1 interactions with the NE, we used single HIV-1 tracking to analyze cores that came into contact with the NE (interactions) and their ability to remain immobile at a single site on the NE (docking) over 8h of virus entry steps. As shown in (Extended. Fig. 11) several of the HIV-1 core tracks detected on the NE corresponded to movement of cores laterally along the NE mask. We therefore analyzed segments of each individual track for docking behavior, as defined by the segments of tracks when a single HIV-1 core remained within a 2-pixel (360 nm) radius for 3 or more frames (>7.5 min). These segments were considered as interactions representative of HIV-1 core docking at the NE. The stringent conditions (2-pixel localization of cores for >3 frames) allowed us to robustly define virus docking and distinguish them from transient interactions along the NE, when cores fail to remain at a given location for more than 5 minutes. The robustness of our docking analysis to track productive docking events was additionally validated by using envelope deficient bald HIV-1 particles (no-VSV), which showed borderline 4.5% docking probability, that was similar to control PF74 and LEN treatments (Fig. 5D), which respectively block interactions with HIV-1 capsids and the nuclear pore complex (NPCs). Compared to these controls, HIV-1 WT and all the MT capsids tested here showed a > 2-fold increase in the fraction of cores that dock at the NE (Fig. 5D), suggesting their ability to effectively interact with NPCs.

The median duration of cores docking at the NE, was similar for WT (~11.8 min), revertant mutants E45A/R132T (~12.7 min), Q63A/Q67E (~12.7 min) and the in-elastic E212/213A mutant (~12.8 min), and significantly higher than the E45A (~8.8 min), Q63A/Q67A (~9.5 min) and E180A (~10.2 min) (Extended Fig. 10). Importantly, the E45A and Q63A/Q67A mutants which fail to enter the nucleus showed similar probabilities for NE-docking as the revertant E45A/R132T and Q63A/Q67E capsid mutants, respectively (Fig. 5D). Moreover, the in-elastic E212/213A capsids which fails to enter the nucleus showed similar level of NE-docking as WT (Fig. 5D and Extended Fig. 10). These results argue that the WT and mutant capsids analyzed here are perfectly capable of engaging the nuclear pore, but elastic properties of cores are additionally required for their passage into the nucleus.

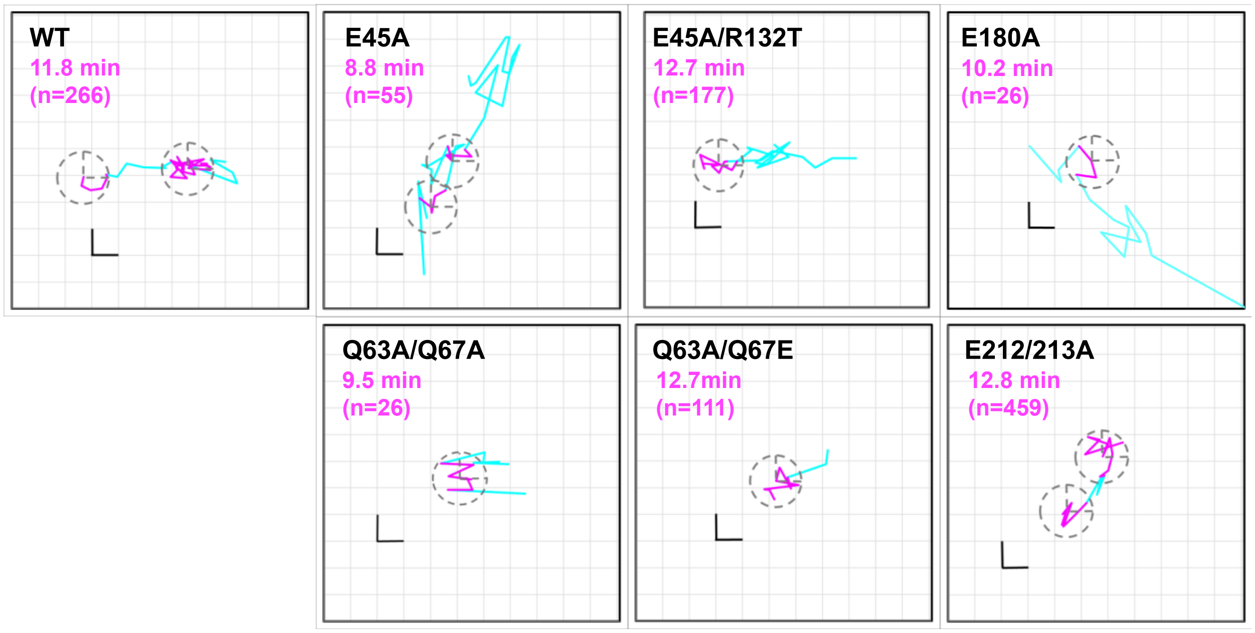


Extended Data Fig. 11 – Single HIV-1 docking analysis for WT and capsid mutants. Single core trajectory along the NE shows local docking (magenta) and lateral movements (cyan). Median docking times (magenta segments, with dashed circles) and number (n) of docking HIV-1 cores analyzed are shown (see also Suppl. Movies 1 – 7). Scale bar 0.36 µm.

1 Francis, A. C. & Melikyan, G. B. Single HIV-1 Imaging Reveals Progression of Infection through CA-Dependent Steps of Docking at the Nuclear Pore, Uncoating, and Nuclear Transport. *Cell Host Microbe* **23**, 536-548 e536, (2018).

2 Burdick, R. C. *et al.* Dynamics and regulation of nuclear import and nuclear movements of HIV-1 complexes. *PLoS Pathog* **13**, e1006570, (2017).
